# Activation of IP10/CXCR3 signaling with highly coincidental with PrP^Sc^ deposit in the brains of scrapie infected mice

**DOI:** 10.1101/2022.04.19.488860

**Authors:** Jia Chen, Cao Chen, Chao Hu, Wei Yang, Lin Wang, Dong-Dong Chen, Yue-Zhang Wu, Qi Shi, Xiao-Ping Dong

## Abstract

Activation of chemokine IP10, also named as CXCL10, and its receptor CXCR3 in CNS is described in some neurodegenerative diseases. Our previous study has also demonstrated an increased brain IP10 levels in several scrapie infected rodent models. However, the detailed alteration of IP10/CXCR3 signaling in CNS during prion infection remains unsettled. Here, we found the increased IP10 signals in the brains of scrapie infected mice mainly localized in the neurons and the activated microglia using various methodologies. The levels of CXCR3 were markedly increased in brains of the scrapie infected mice and in the prion infected cell line SMB-S15. The increased CXCR3 mainly distributed in neurons. Obviously morphological colocalizations of PrP/PrP^Sc^ with IP10 and CXCR3 in the brains of scrapie infected mice were observed in the assays of immunohistochemistry (IHC) and immunofluorescence. Additionally, IHC analysis with whole brain sections demonstrated that the increased IP10 and CXCR3 accumulated in the brain regions with more PrP^Sc^ deposits. Co-immunoprecipitation and biomolecular interaction assays identified the evidence for the molecular interactions of PrP with IP10 and CXCR3. Compared to the normal partner cell line SMB-PS, the more portion of IP10 accumulated insides of prion infected SMB-S15 cells. Removal of prion replication in SMB-S15 cells by resveratrol converted the pattern of the accumulation and secretion of cellular IP10. Our data here demonstrate an activation of IP10/CXCR3 signaling in the brain tissues of prion infection, highly coincidental with PrP^Sc^ deposit. Modulation of brain IP10/CXCR3 signaling is potential therapeutic target for reducing the progression of prion diseases.

## Introduction

Prion disease, also known as transmissible spongiform encephalopathies (TSEs), is a type of fatal degenerative disease that can invade the central nervous system of humans and many mammals [1, 2]. The causative agent is prion, a self-replicating protein without nucleic acid. Current knowledge believes that the central conception of prion is that the conformational conversion from a cell-type prion protein (PrP^C^) normally expressed in central nerve system (CNS) and other tissues to the infectious and pathogenic scrapie-like prion protein (PrP^Sc^), due to spontaneous alteration, genetic mutations and exogenous infection. In addition to the wide deposits of PrP^Sc^, loss of neurons and reactive proliferation of glial cells in CNS, activation of innate or unspecific immunity, e.g., activation of microglia and complement system as well as increases of various cytokines and chemokines, are also addressed [3].

The innate immune system is a sort of host defense response aiming to eliminate the invaded infectious agents. As important parts of the innate immune system, activations of glial cells and various inflammatory factors in the brain tissues are frequent observed both in acute and chronic CNS infectious diseases. Innate immune system in CNS is also activated in the patients suffered from many neurodegenerative diseases, which is closely associated with aberrant deposits of aggregated and misfolded proteins, such as amyloid-β in Alzheimer disease (AD), α-synuclein in Parkinson disease (PD), and prions in prion disease [4]. Abnormal upregulations of dozens of cytokines are described in the brains of various scrapie-infected experimental mouse models, as well as in the cerebral spinal fluid (CSF) and brains of Creutzfeldt-Jacob disease (CJD) patients, including chemokine (C-X-C motif) ligand 10 (CXCL10) that is also named as interferon gamma-induced protein 10 (IP10) [5–11].

Chemokines are a group of small molecular proteins ranging from 8 to 10 kDa. According to the arrangement of cysteines in N-terminus, chemokines can be divided into four main subfamilies, CXC, CC, CX3C and XC subfamilies [12, 13]. IP10 is a secretory protein with a molecular weight of 10 kDa belonging to CXC subfamily. IP10 induces the chemotactic response of monocytes and macrophages, as well as other immune cells, such as T cells, NK cells and DC cells [14]. IP10 also involve in many physiological activities, such as white blood cell transport, acquired immune response, inflammation, hematopoiesis, and angiogenesis [15–21]. The biofunction of IP10 is mediated via binding to the chemokine receptor CXCR3 [22]. CXCR3 is a membrane protein with 7 transmembrane units and belongs to G protein-coupled receptor (GPCR). The main ligands of CXCR3 include CXCL9, CXCL10 and CXCL11 [14, 23, 24]. Using Luminex assay and ELISA, we have found that the IP10 levels in the brain tissues of mice infected with different scrapie agents were abnormally increased, meanwhile that of IP10 was slightly increased in the cerebrospinal fluid (CSF) of the patients with sporadic Creutzfeldt-Jakob disease [25]. However, the morphological distribution of IP10 and the potential alteration of CRCX3 in the brains of prion diseases remain unclear.

In the present study, we found that the increased IP10 signals in the brain tissues of scrapie infected mice mainly localized in the neurons and the activated microglia. The levels of CXCR3 were markedly increased in the brains of the mice infected with scrapie agents and in the prion infected cell line SMB-S15. The increased CXCR3 mainly distributed in neurons. Obvious colocalizations of PrP/PrP^Sc^ with IP10 and CXCR3 in the brain tissues of scrapie infected mice were observed. In addition, the increased IP10 and CXCR3 accumulated in the brain regions with more PrP^Sc^ deposits. Using coimmunoprecipitation assays and biomolecular interaction analysis system, evidence for the molecular interactions of PrP with IP10 and CXCR3 molecules was identified. Compared to the normal partner cell line SMB-PS, the increased IP10 in SMB-S15 cells accumulated more insides of cells. Removal of prion replication in SMB-S15 cells by resveratrol converted the accumulation and secretion of cellular IP10.

## Result

### IP10 localizes at neurons and microglia, but not at astrocytes in the brain tissues of scrapie-infected mice

Using the techniques of Luminex and ELISA, the upregulation of brain IP10 in the scrapie infected experimental rodents has been addressed [25]. In this study, the levels of brain IP10 in mice infected with scrapie agents 139A and ME7 at terminal stage were further analyzed with IP10-specific Western blots. Stronger IP10 signals were detected in the group of scrapie infected mice compared to that of age-matched normal ones (Fig. 1A). To identify the expression and localization of IP10 during scrapie infection among the different types of cells in the CNS morphologically, the sections of cortex regions from 139A- and ME7-infected mice were double-stained using immunofluorescence with NeuN, GFAP or Iba1 specific antibodies, respectively. Obviously more IP10-specific signals (green) were observed in the preparations of 139A- and ME7-infected mice at terminal stage (Fig. 1B-D). Double staining of the tissues with the antibodies for IP10 and biomarkers of the various cell types revealed that the IP10-positive signals (green) colocalized with the NeuN-positive cells (red) (Fig. 1B) and Iba1-positive cells (red) (Fig. 1C) in the cortex region of scrapie-infected mice, but seemed not to overlap with the GFAP-positively stained cells (Fig. 1D). It highlights that the increased IP10 signals in the brain tissues of scrapie infected mice mainly localize at the neurons and the activated microglia.

**Figure 1.**
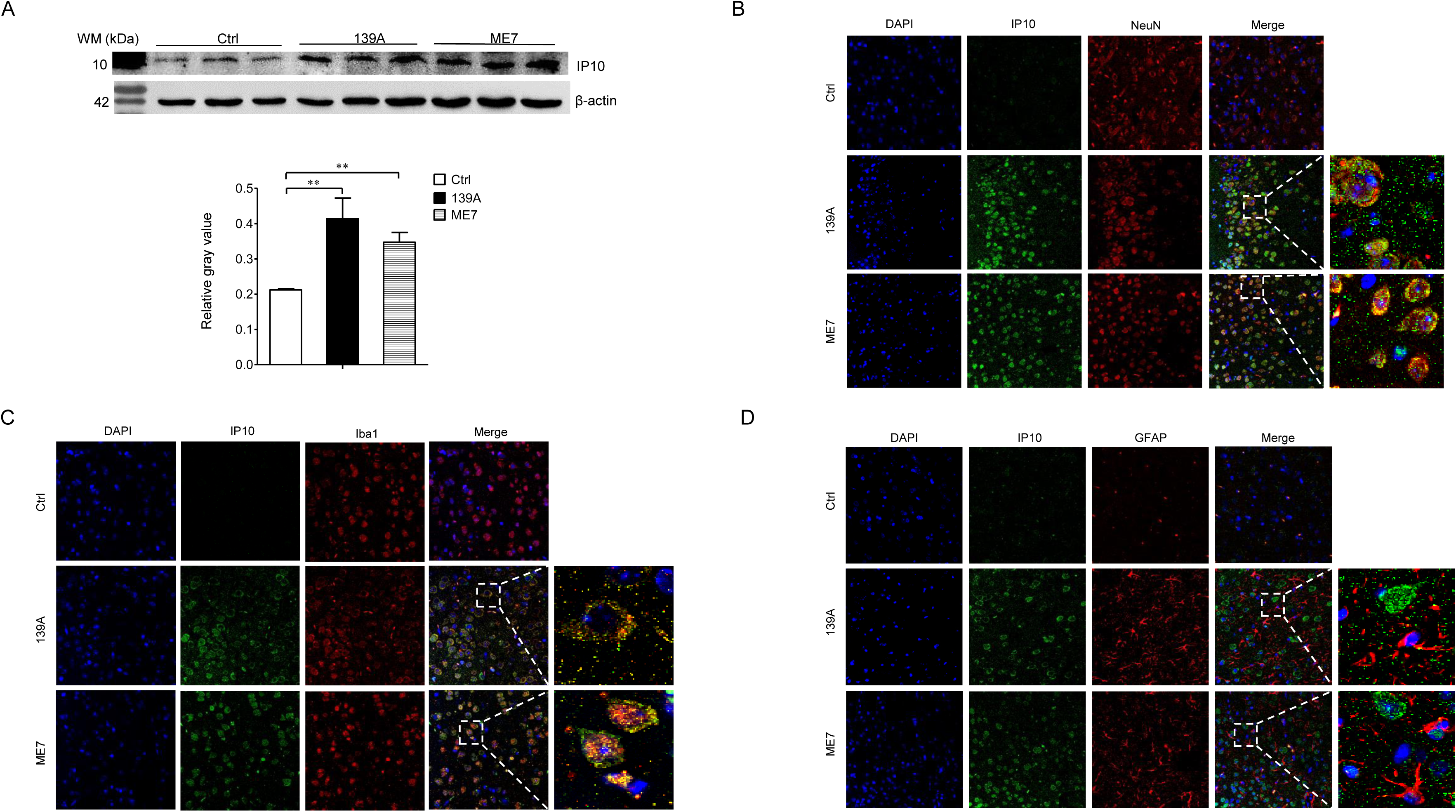
Increase of IP10 in the brains of scrapie agents infected mice. **A.** Western blot of IP10 in the brain homogenates from agents 139A and ME7 at terminal stage (n=3). Equal amounts of brain homogenates of normal and infected mice were loaded into 12% SDS-PAGE. β-actin was used as an internal control. Molecular weight is marked on the left and various specific immunoblots are marked on the right. Quantitative assays determined by densitometry relative to IP10/β-actin are showed on the bottom. Graphical data denote mean + SEM (n=3). Statistical differences compared with controls are illustrated on the top. **B.** Representative images of double stained of IP10 (green) and NeuN (red) in the brain sections of 139A and ME7 infected mice. **C.** Representative images of double stained of IP10 (green) and Iba1 (red) in the brain sections of 139A and ME7 infected mice. **D.** Representative images of double stained of IP10 (green) and GFAP (red) in the brain sections of 139A and ME7 infected mice. The enlarged images are showed on the right.

### Upregulation of IP10 in prion-infected cell line SMB-S15

To explore the possible change of IP10 in the cultured cells with prion replication, the expression of IP10 in SMB-15 cells and its normal partner cell line SMB-PS were evaluated. Quantitative RT-PCR assays with IP10-specific primers revealed an increased transcription of the specific mRNAs in SMB-S15 cells, showing statistical difference to SMB-PS cells (Fig. 2A). Western blot with anti-IP10 identified the specific bands in the lysates of SMB-15 cells but not in that of SMB-PS cells (Fig. 2B). Measurement of IP10 in the cellular lysates with a commercial ELISA kit also proposed significantly higher expressing levels in SMB-S15 cells (Fig. 2C). Furthermore, markedly stronger IP10 signals (green) were observed in SMB-S15 cells, showing significantly higher average IOD values compared to SMB-PS cells, either in the fractions of cytoplasm or cell nucleus (Fig. 2D). It implies a higher endogenous expression of IP10 in the cell line with persistent replication of prions.

**Figure 2.**
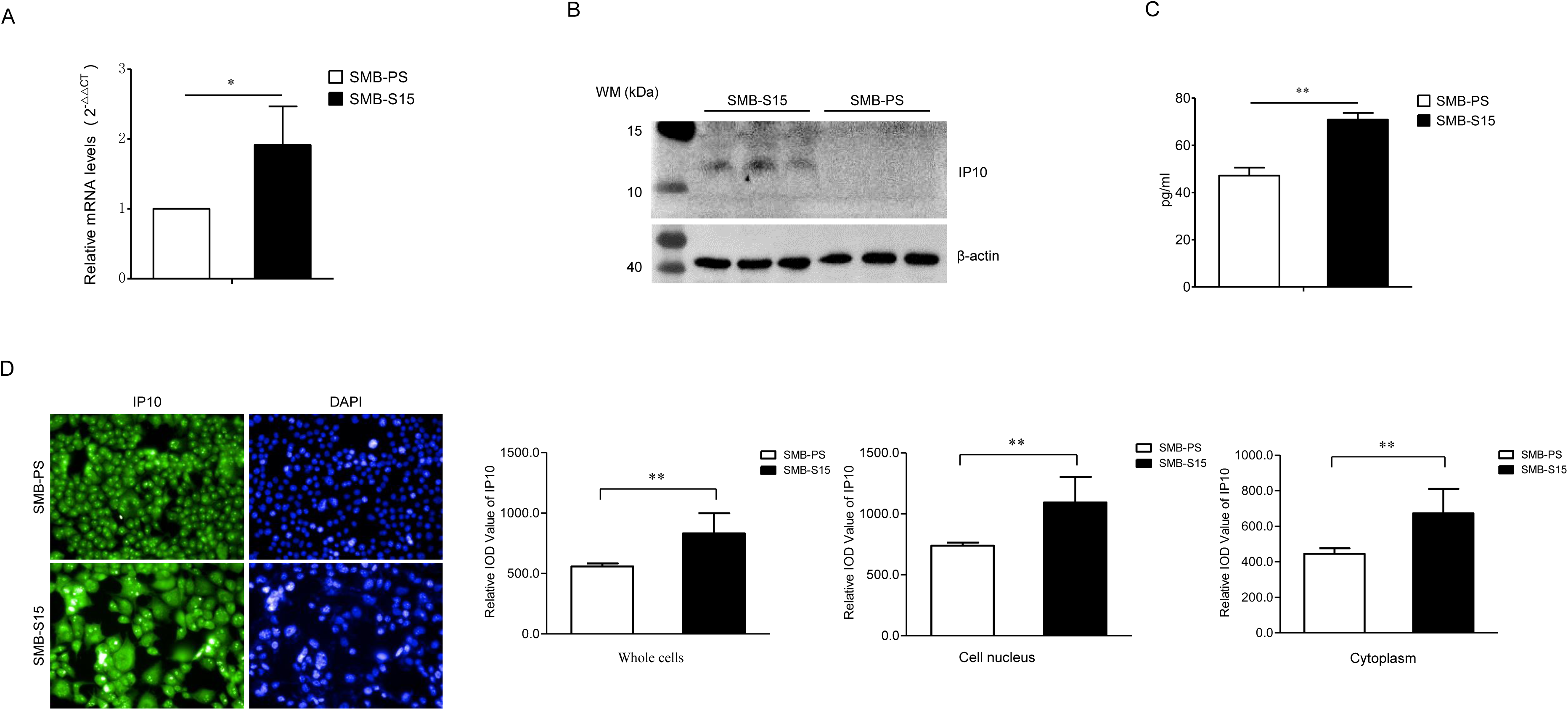
Increase of IP10 in prion infected cell line SMB-S15. **A.** IP10 specific qRT-PCR. Total RNAs from three batches of cultured SMB-S15 and SMB-PS cells were analyzed for the transcriptional levels of IP10 by the established IP10 qRT-PCR. Y-axis represents the values of 2^-ρρCT^ of SMB-S15 and SMB-PS cells. **B.** Western blot of IP10 in three batches of each SMB cells. **C.** Measurement of IP10 level with ELISA kit. The amounts of IP10 (pg/ml) are indicated on Y-axis. **D.** Representative images of IP10 specific IFAs. The intensity optical density (IOD) values of IP10 in the fractions of whole cells, cytoplasm and nucleus of two SMB cell lines in three randomly selected fields are automatically calculated with the software image J separately.

### Upregulation of CXCR3 in the prion infected cells and in the brains of scrapie-infected mice collected at the moribund stage

To evaluate the potential change of CXCR3 during prion infection, the lysates of the SMB-PS and SMB-S15 cells were enrolled in IP10-specific Western blots. The level of CXCR3 in SMB-S15 were higher than that of SMB-PS cells, with statistical difference in quantitative assay (Fig. 3A). CXCR3-specific IFA illustrated stronger green signals in SMB-S15 than that in SMB-PS cells, that the IOD value of CXCR3 in SMB-S15 cells was significantly higher than that of SMB-PS cells after normalized with the individual data of nucleus in the quantitative analysis (p<0.01, Fig. 3B), reflecting a relatively higher CXCR3 level in prion infected cells.

**Figure 3.**
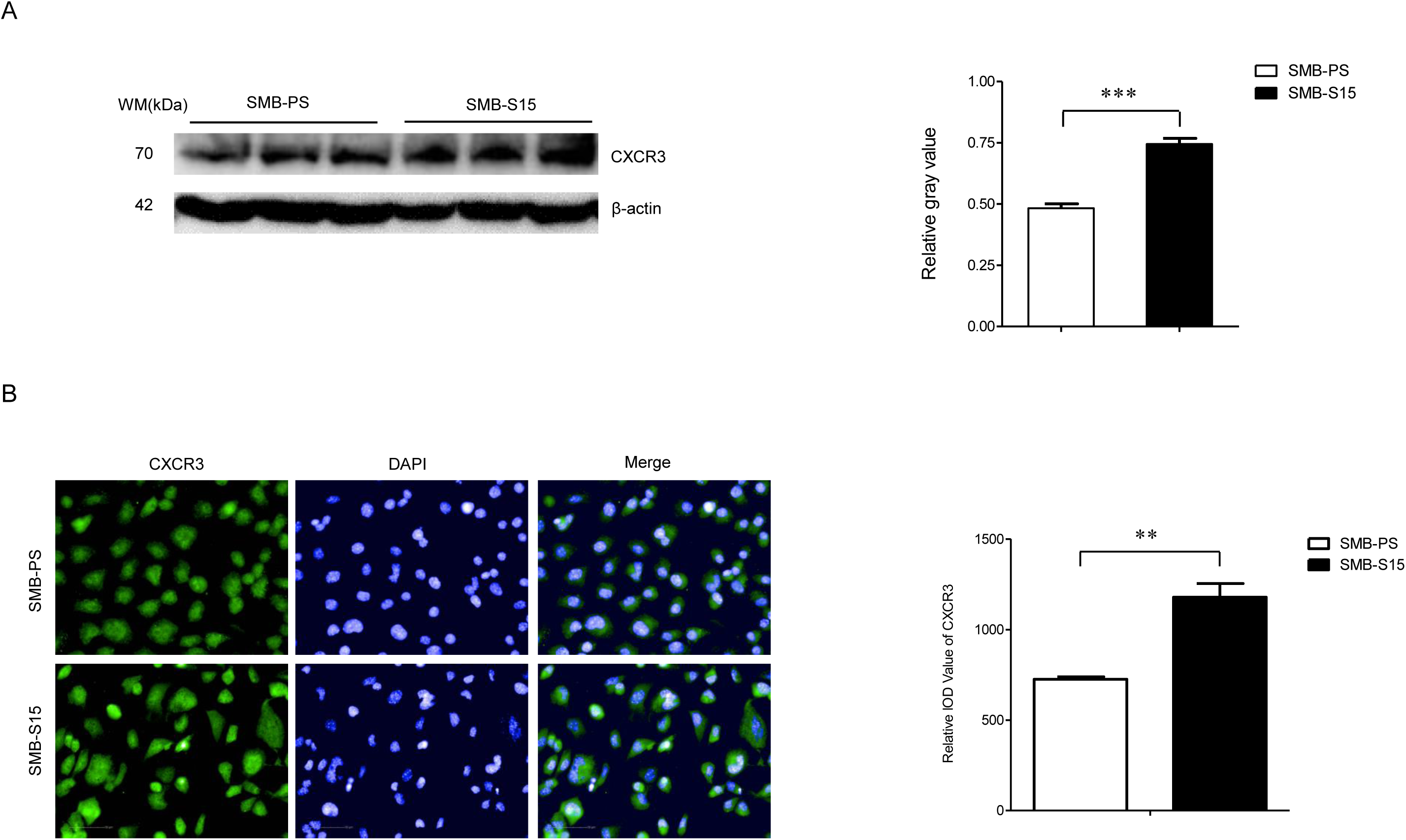
Increase of CXCR3 in prion infected cell line SMB-S15. **A.** Western blot of IP10 in three batches of each SMB cells. Quantitative assays are determined by densitometry relative to IP10/β-actin and graphical data denote mean + SEM (n=3). **B.** CXCR3 specific IFAs. The IOD values of IP10 in SMB-S15 and SMB-PS cells are automatically calculated with the software image J.

To evaluate the alterations of CXCR3 in the brain tissues of experimental rodents infected with scrapie agents, the brain samples from 139A- and ME7-infected mice (n=3) were enrolled, respectively. The transcriptional status of CXCR3 in brains of scrapie-infected mice were assessed by real time PCR with CXCR3-specific primer pairs, using the housekeeping gene encoding β-actin as the internal control. After being equilibrated with the Ct value of respective β-actin, the 2^−ΔΔCt^ values of CXCR3 were calculated from the data of three independent reactions. The levels of CXCR3 transcripts in the brains of 139A- and ME7-infected mice were higher than that of controls, showing statistical difference compared to the controls (Fig. 4A).

**Figure 4.**
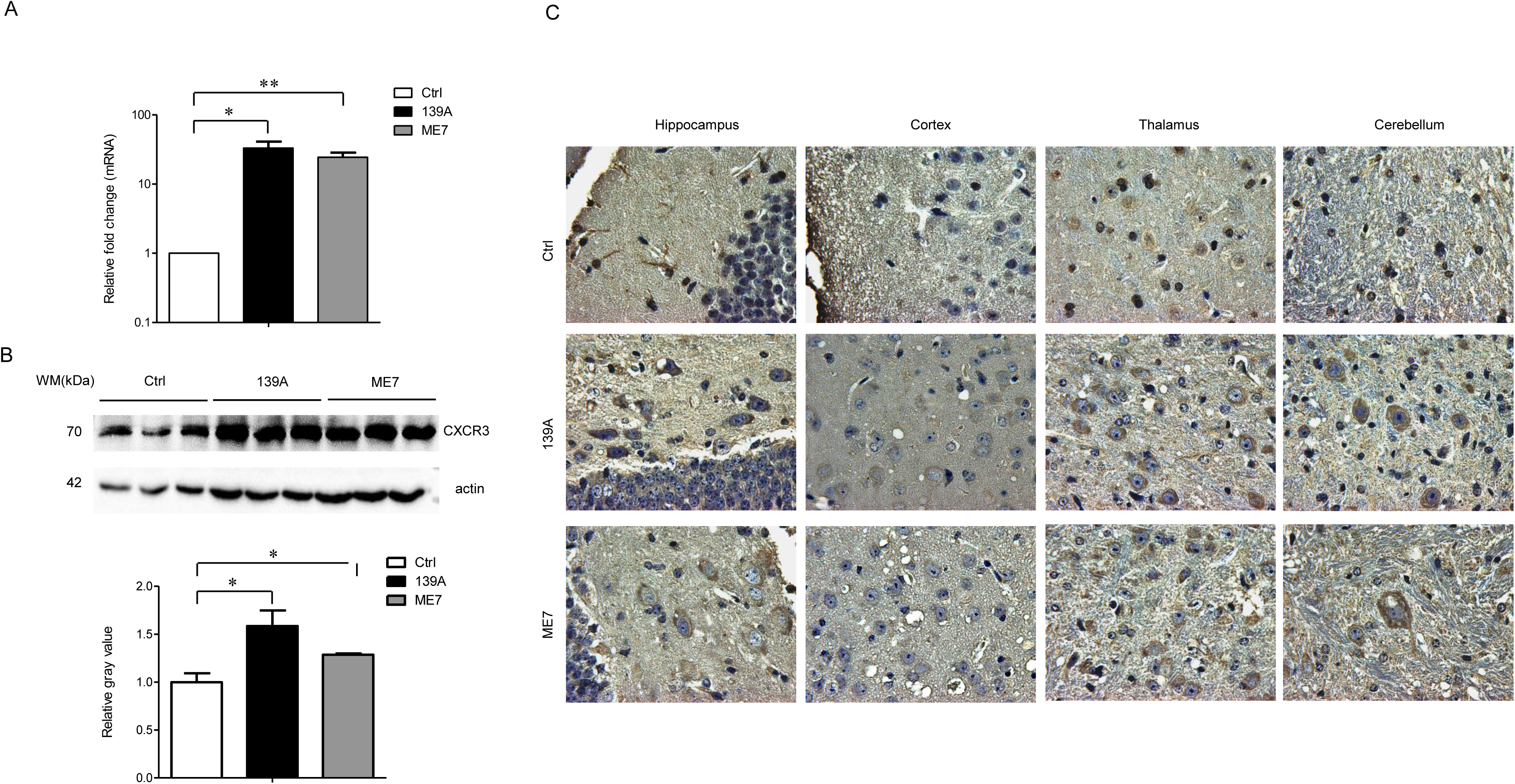
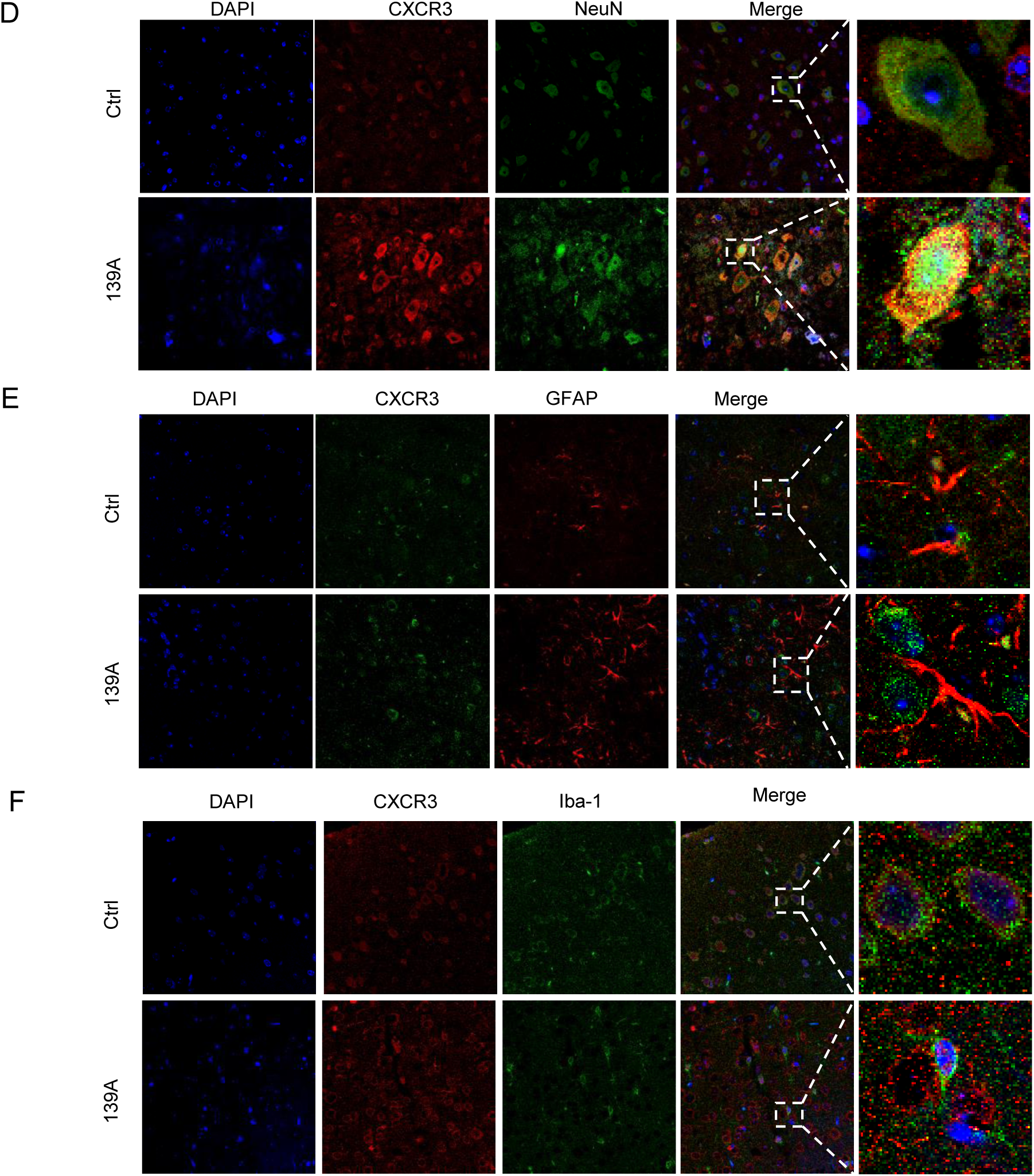
Increase of CXCR3 in the brains of scrapie agents infected mice. **A.** IP10 specific qRT-PCR. The extracted total RNAs of normal and 139A, ME7 infected mice (n=3 each) were subjected into IP10 qRT-PCR. Y-axis represents the values of 2^-ρρCT^. **B.** CXCR3 specific Western blot. Equal amounts of brain homogenates of normal and infected mice (n=3 each) were subjected into IP10 specific Western blot. Quantitative assays are determined by densitometry relative to IP10/β-actin and graphical data denote mean + SEM. **C.** Representive graphs of CXCR3 specific IHC assays of the brain sections of normal, 139A and ME7 infected mice. The brain regions of hippocampus, cortex, thalamus, and cerebellum are indicated on the tops of images. **D.** Representative images of double stained of IP10 (green) and NeuN (red) in the brain sections of normal and 139A infected mice. **E.** Representative images of double stained of IP10 (red) and Iba1 (green) in the brain sections of normal and 139A infected mice. **F.** Representative images of double stained of IP10 (green) and GFAP (red) in the brain sections of normal and 139A infected mice. The enlarged images are showed on the right.

Western blots showed stronger CXCR3 signals in the brains of the 139A- and ME7-infected mice compared to the age matched healthy controls, with statistical difference in the quantitative assays (Fig. 4B). To add further insight into the expressions of CXCR3 in various brain regions, the sections of four different brain regions of 139A- and ME7-infected and age-matched healthy mice were subjected to CXCR3-specific IHC assays, including hippocampus, cortex, thalamus and cerebellum. Large amounts of the CXCR3-specific dark brown staining were observed in all tested brain sections of 139A- and ME7-infected mice compared with those of the control group, majority locating within the cell bodies (Fig. 4C). Subsequently, the brain sections of 139A-infected and normal mice employed into double stained IFAs in order to see the CXCR3 distributions on different types of cells in brain tissues. More CXCR3 signals were observed in the brain slices of 139A-infected mice than that of normal control (Fig. 4D-F). Analyses of the merged images identified colocalized signals (yellow) of CXCR3 with NeuN-positive cells (Fig. 4D), but not with GFAP- (Fig. 4E) and Iba1- (Fig. 4F) stained cells. Those data indicate that the CXCR3 levels were upregulated in the brains during prion infection and the increased CXCR3 seems to distribute mainly in neurons.

### Morphological associations between PrP^Sc^, IP10 and CXCR3 in the brain tissues of scrapie infected mice

To evaluate the morphological colocalization of increased CXCR3 with PrP, the brain sections of normal and 139A-infected mice were double stained immunofluorescently with anti-CXCR3 and anti-PrP. On the background of the increased CXCR3 and PrP signals, notably more colocalized signals (yellow) insides of cellular structures were identified in the merged images of the different brain regions of 139A-infected mice, such as cortex (Fig. 5A), hippocampus (Fig. 5B) and cerebellum (Fig. 5C), whereas less but repeatedly observable colocalized signals were also detected in the brain tissues of normal animals.

**Figure 5.**
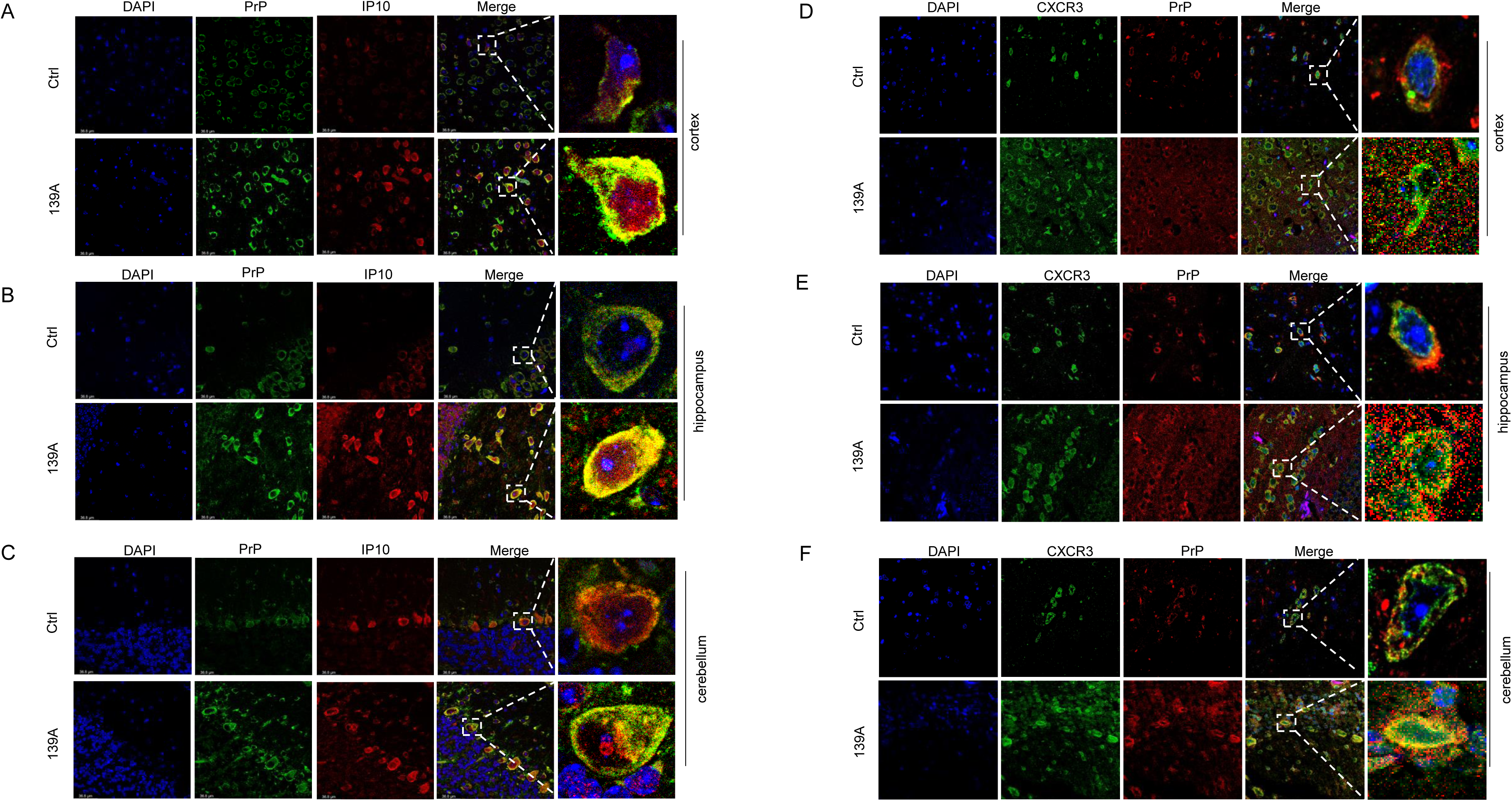
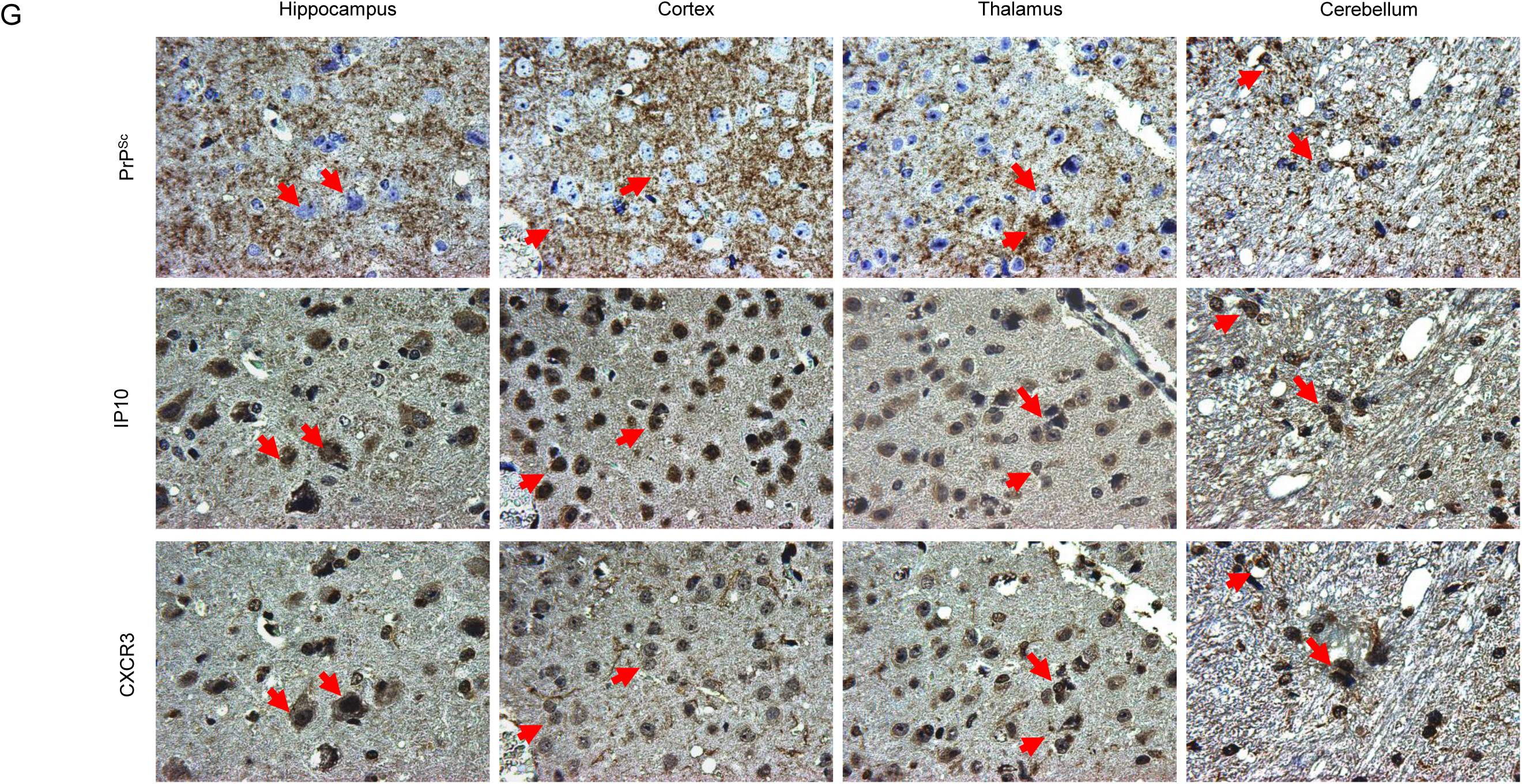
Morphological colocalizations among PrP/PrP^Sc^, IP10 and CXCR3 in the brains of scrapie infected mice. **A-C.** Representative images of double stained IFA of PrP (green) and IP10 (red) in the regions of cortex (A), hippocampus (B), and cerebellum (C) of normal and 139A infected mice. **D-F.** Representative images of double stained IFA of PrP (red) and CXCR3 (green) in the regions of cortex (D), hippocampus (E), and cerebellum (F) of normal and 139A infected mice. The enlarged images are showed on the right. **G.** Representative IHC images of PrP^Sc^, IP10 and CXCR3 in the serial brain sections of 139A infected mice. For PrP^Sc^, the brain sections were treated with GdnHCl and subsequently stained with PrP specific mAb prior to IHC assay. Arrows indicate the colocalized signals.

To determine the morphological association between PrP^Sc^, IP10 and CXCR3 in the brains with prion infection, the serial brain sections of 139A-infected mice were prepared. The IHC assays for IP10 and CXCR3 were conducted routinely, while that for PrP^Sc^ was performed after removal of normal PrP by treatment with GdnHCl. As shown in Figure 5D, large amounts of brown signals deposited at the same positions in the serial section stained by anti-IP10, anti-CXCR3 and anti-PrP, respectively. Co-deposits of PrP^Sc^, IP10 and CXCR3 distributed widely in various brain regions, including cortex, hippocampus, thalamus and cerebellum (Fig. 5D).

Furthermore, the whole brain sections of 139A-infected mice were immunohistochemically stained with anti-IP10, anti-CXCR3 and anti-PrP, respectively. With the help of the software Imagescope equipped in Leica Aperio CS2, the percent positive of PrP^Sc^, IP10 and CXCR3 in the regions of hippocampus, cerebellum, cortex and thalamus were automatically scanned and calculated. As illustrated in Figure 6, in hippocampus, cerebellum, cortex and thalamus the percent positive of PrP^Sc^ (Fig. 6A) were 6.54%, 22.99%, 21.51% and 6.57%, while those of IP10 (Fig. 6B) and CXCR3 (Fig. 6C) were 9.56%, 25.49%, 26.09% and 11.92%, 7.88%, 27.30%, 17.12% and 10.07%, respectively. Remarkably, more positive signals of those three biomarkers deposited in the regions of cortex and cerebellum. It highlights that the increased IP10 and CXCR3 accumulated in the brain regions with more PrP^Sc^ deposits.

**Figure 6.**
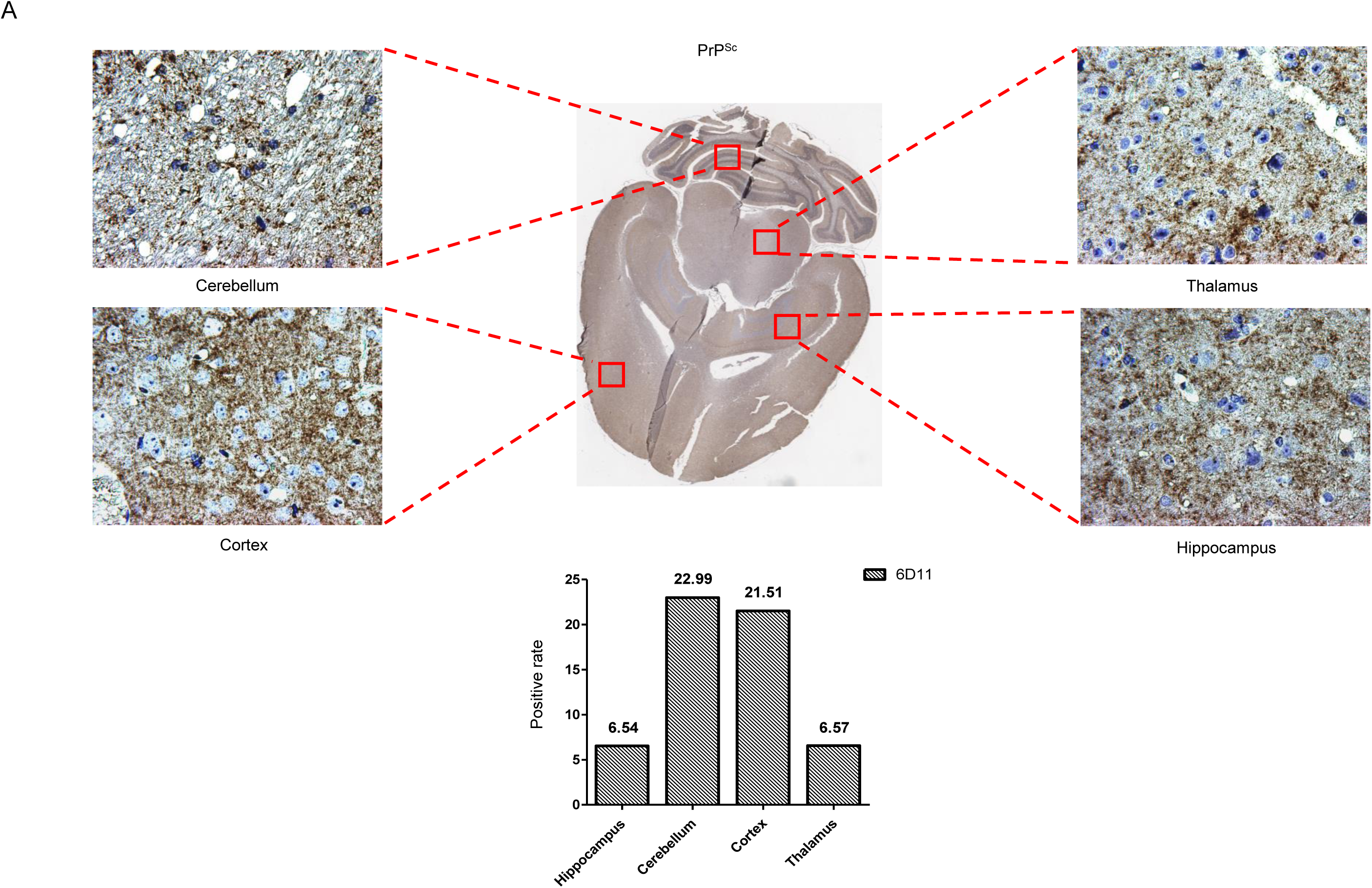
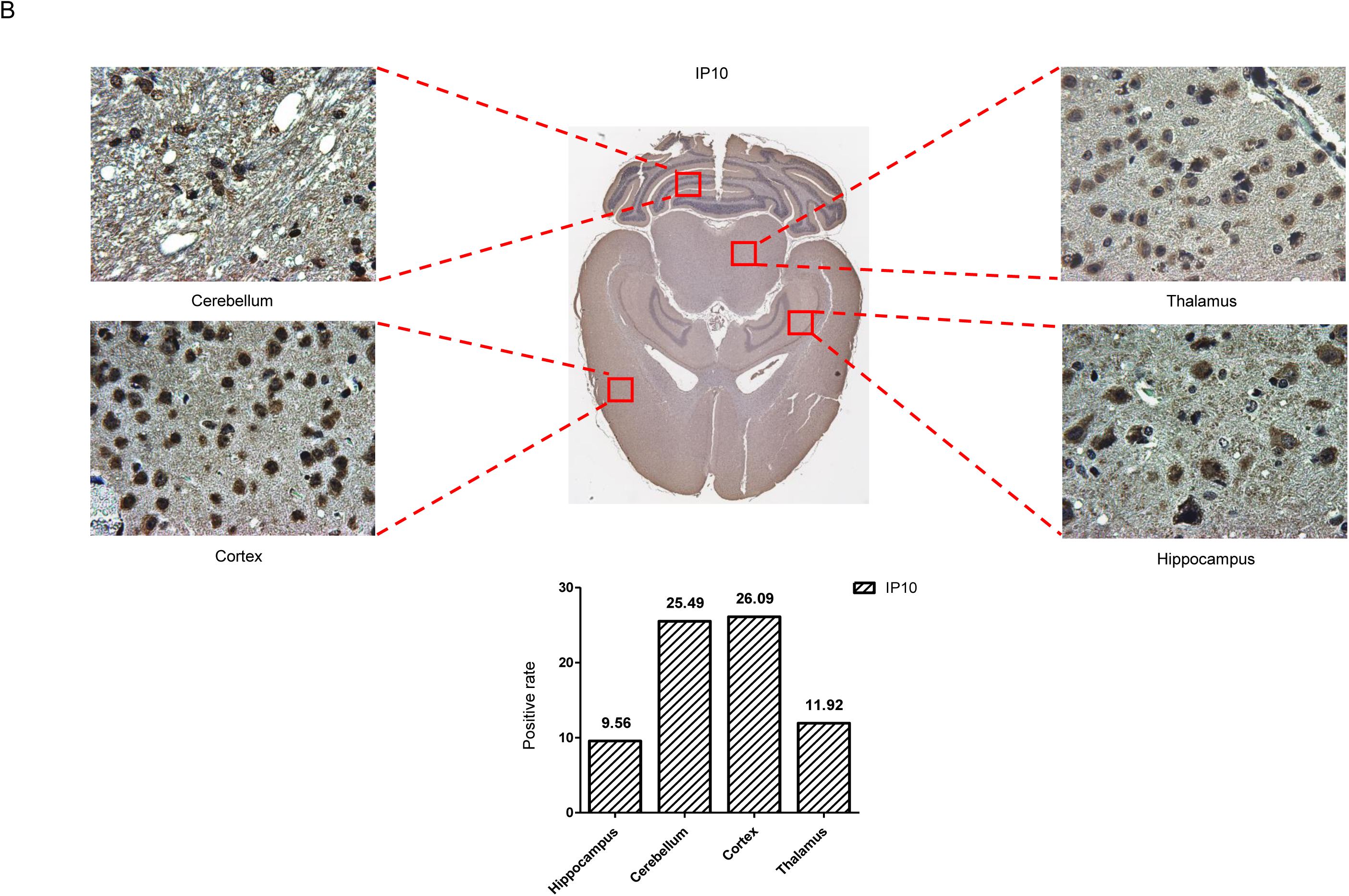
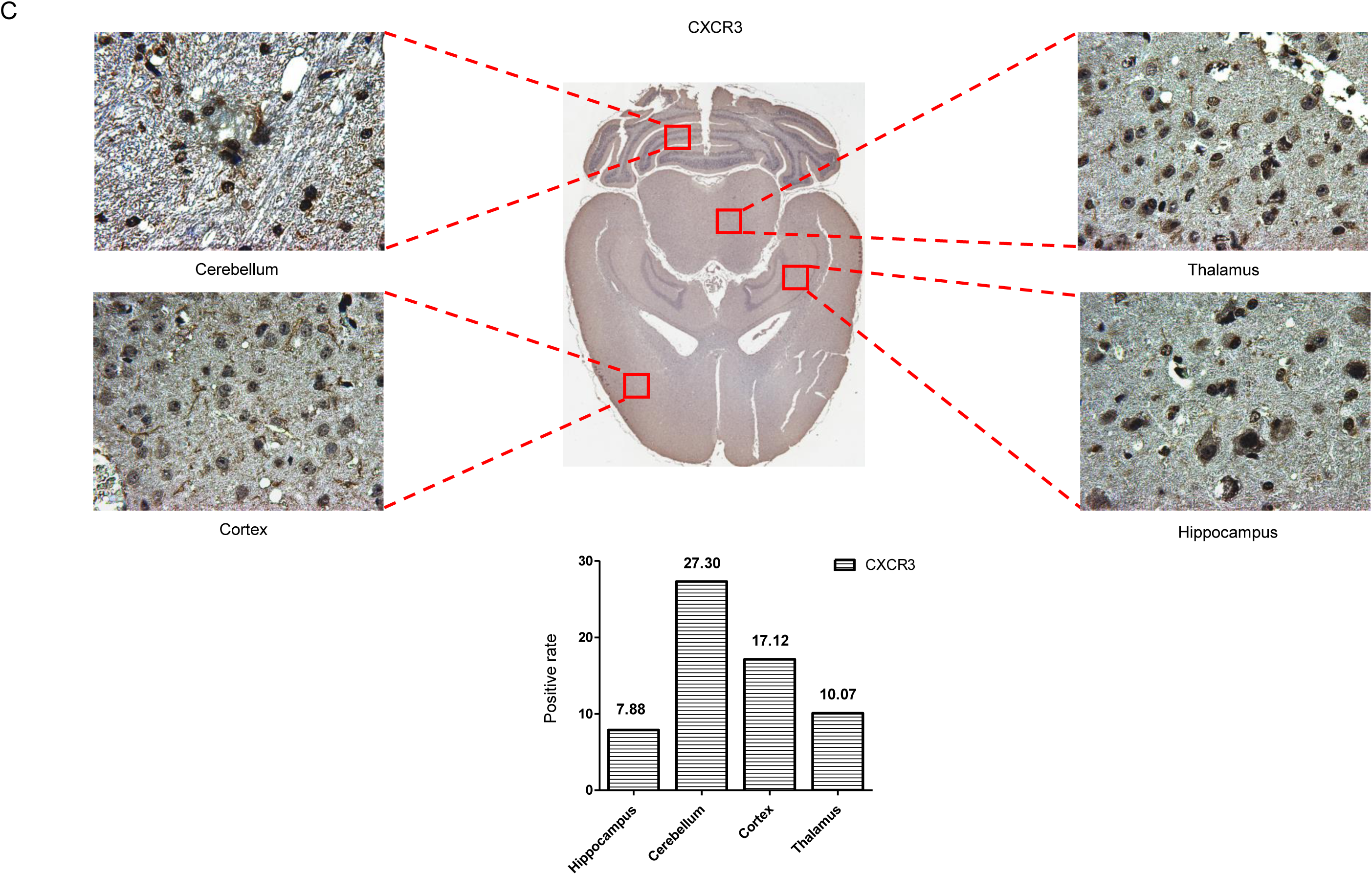
Co-distributions of PrP^Sc^, IP10 and CXCR3 in the different brain regions of scrapie infected mice. Whole brain sections of 139A infected mice were separately stained by PrP^Sc^, IP10, CXCR3 specific mAbs in IHC assays. **A.** PrP^Sc^. **B.** IP10. **C.** CXCR3. The whole brain IHC pictures generated by software Imagescope equipped in Leica Aperio CS2 are indicated in the central and the representive graphs of the regions of hippocampus, cortex, thalamus, and cerebellum are showed around. The positive percentages of PrP^Sc^, IP10 and CXCR3 are illustrated below.

### Molecular interactions between PrP, IP10 and CXCR3

Molecular interaction between IP10 and CXCR3 has been addressed elsewhere . To address the potential molecular binding between PrP and IP10, and between PrP and CXCR3, the lysates of SMB-S15 cells were subjected into the immunoprecipitation assays using anti-IP10 or anti-CXCR3 as the capturing antibody. and anti-PrP as the detecting one. Prior to the Western blots, the precipitated products were digested with 20 mg/ml PK. Clear PK-resistant PrP bands were detected in the preparations captured by either anti-IP10 (Fig. 7A) or anti-CXCR3 (Fig. 7B), but not in the reactions precipitated with isotype mouse IgG. It indicates that the PrP, particularly PrP^Sc^, in SMB-S15 cells can form complexes with both IP10 and CXCR3. To obtain more evidence, prokaryotic recombinant full-length human PrP (rHuPrP23-231) was expressed and purified, and its molecular interaction with a commercially supplied recombinant human IP10 protein was measured with ForteBio Octet RED96E biomolecular interaction analysis system. As shown in Figure 7C, a notable binding activity was observed between the immobilized rHuPrP23-231 and the input IP10 with a *K_D_* value of 0.4739 nM, which provides definite evidence for the interaction of PrP with IP10 molecules.

**Figure 7.**
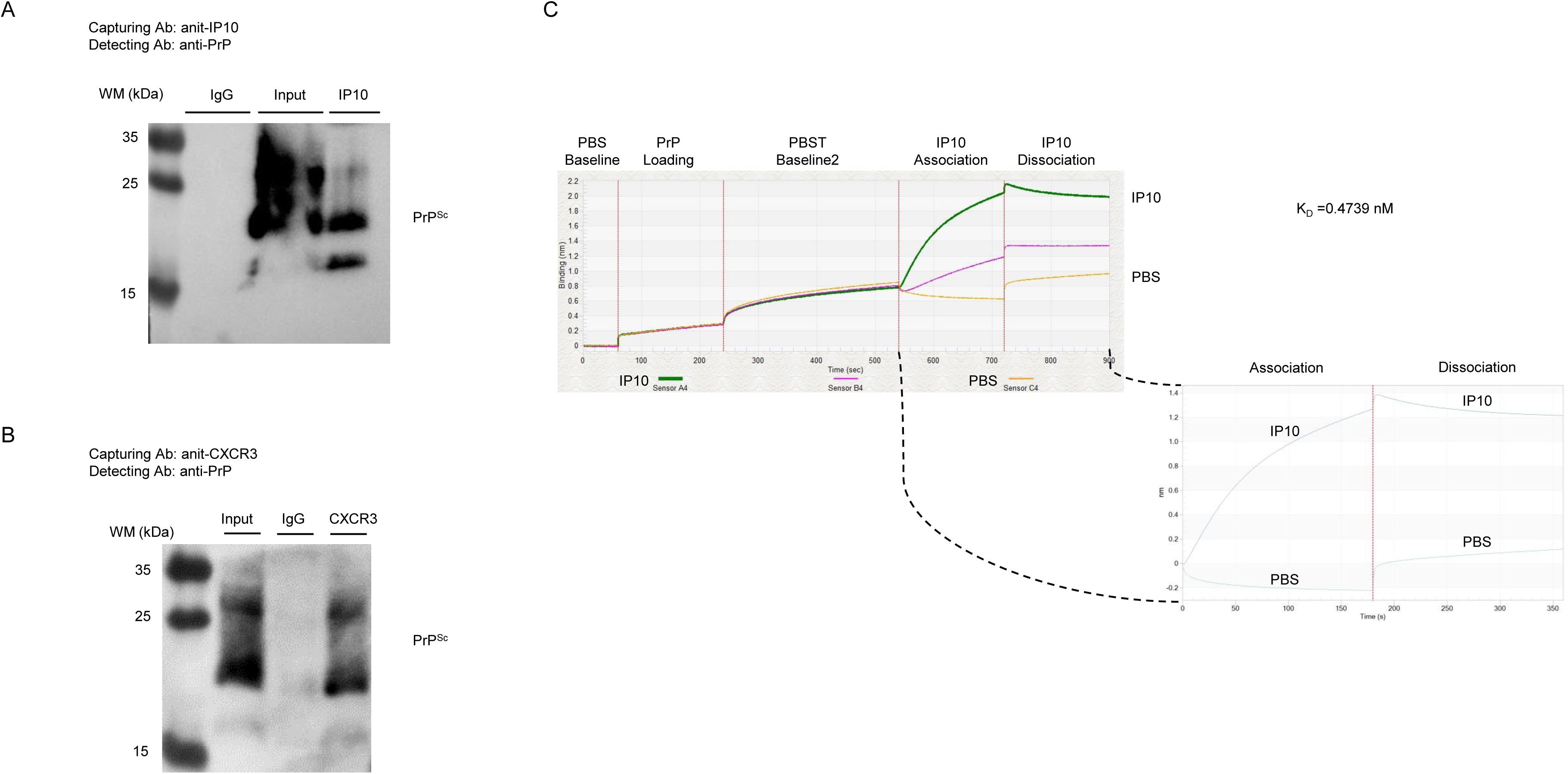
Molecular interactions between PrP/PrP^Sc^ and IP10, PrP/PrP^Sc^ and CXCR3. **A.** Co-IP for PrP^Sc^ and IP10. The brain homogenate of 139A infected mice was captured with anti-IP10 or isotypic IgG, and blotted with anti-PrP. Prior to SDS-PAGE, the eluted product precipitated by anti-PrP was digested with 20 mg/ml PK. **B.** Co-IP for PrP^Sc^ and CXCR3. The brain homogenate of 139A infected mice was captured with anti-CXCR3 or isotypic IgG, and blotted with anti-PrP. Prior to SDS-PAGE, the eluted product precipitated by anti-PrP was digested with 20 mg/ml PK. **C.** Measurement of binding affinities between recombinant full-length human PrP and IP10 using Biolayer Interferometry. The K_D_ (M) value between PrP and IP10 is automatically generated.

### Accumulation of increased IP10 insides of SMB-S15 cells and reversion of the accumulation of IP10 and increased CXCR3 via removal of cellular PrP^Sc^ by resveratrol

To address the distribution of the increased IP10 insides and outsides of SMB-S15 and SMB-PS cells, the cultured mediums and the cells were collected 24, 48 and 72 h post-passage separately and the levels of IP10 in those two fractions were measured by the commercial IP10 ELISA kit. Interestingly, at all three time-points the amounts of IP10 insides of SMB-S15 cells were higher than that of SMB-PS cells, whereas the amounts of IP10 in the cultured medium of SMB-S15 cells were significantly lower than that of SMB-PS cells (Fig. 8A), highlighting the increased IP10 in the prion infected cells distributing mainly insides of the cells. To compare the activity of the IP10 in the medium fractions of two SMB cell lines, SMB cells were co-cultured with a microglia cell line BV2 separated with semipermeable membrane. After stained the membranes with crystal violet, the migrated BV cells were counted. As expected, more numbers of migrated BV2 cells were identified in the membranes in the co-culture preparations with SMB-PS cells compared to SMB-S15 cells (Fig. 8B).

**Figure 8.**
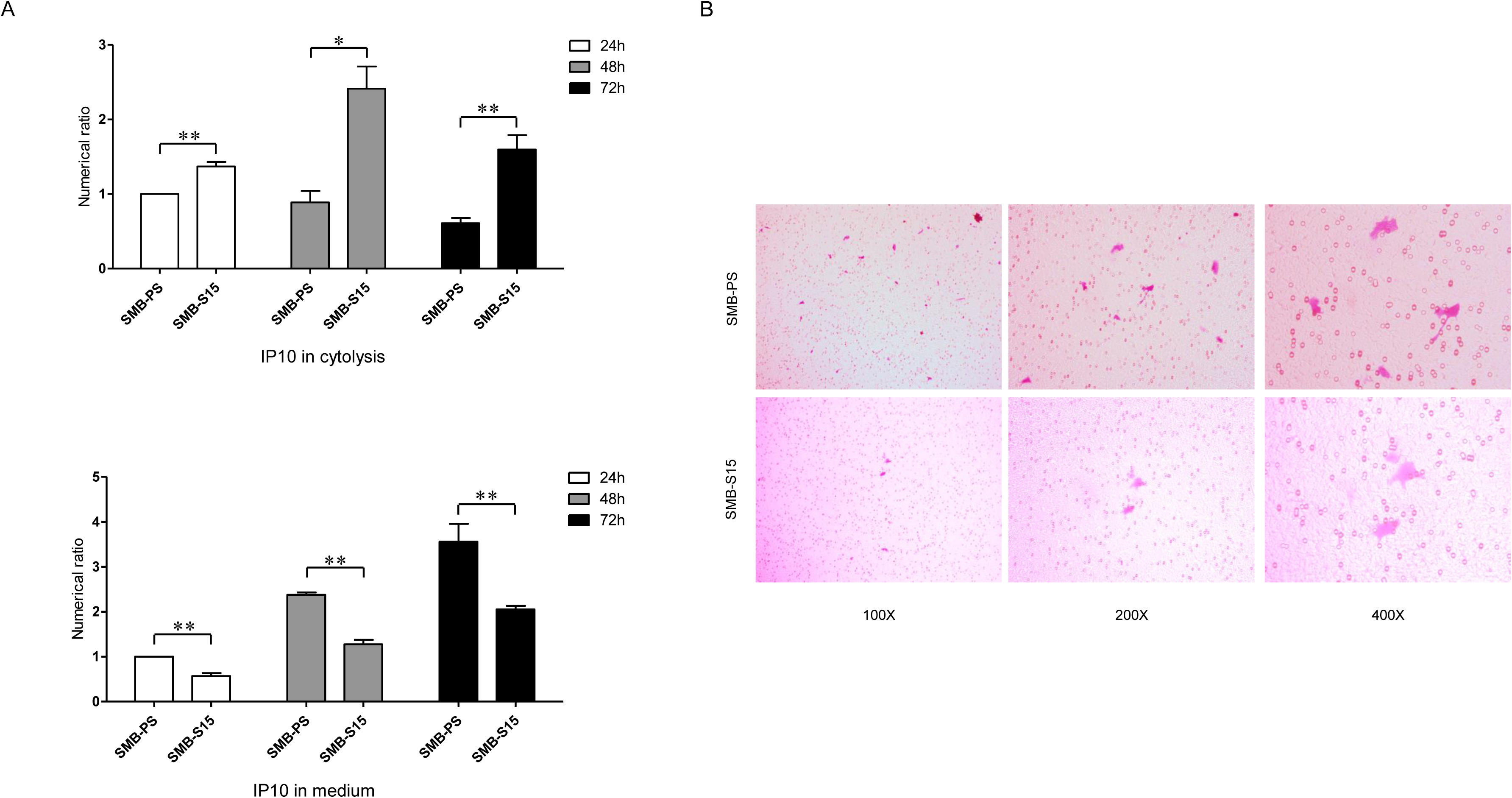
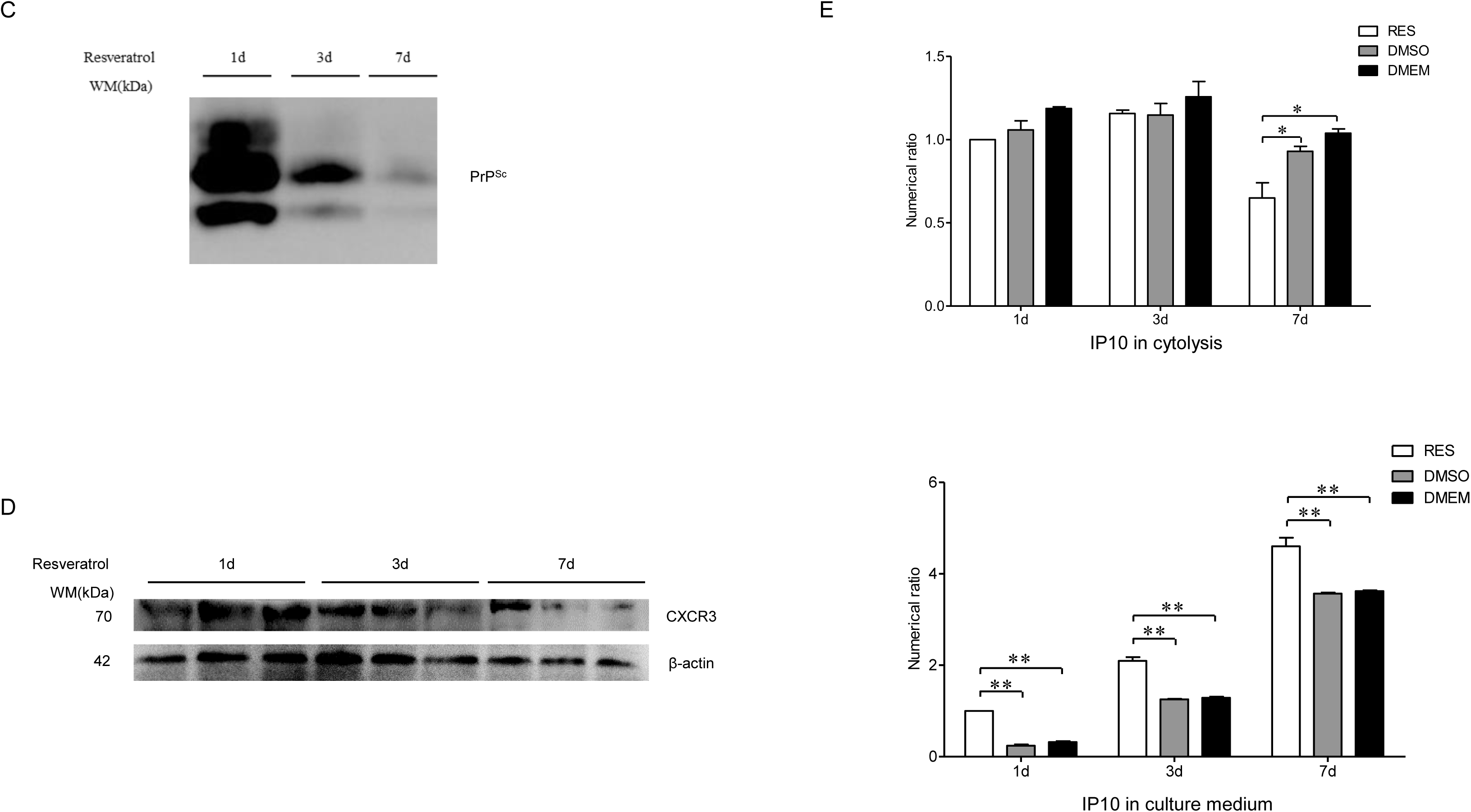
Distributions of IP10 and CXCR3 in the fractions of cellular lysates and cultured medium before and after treatment of resveratrol. **A.** Measurement of IP10 in the cytolysis and medium of SMB-S15 and SMB-PS cells with ELISA before treatment of resveratrol. Cells were collected 24, 48 and 72 h after passaged and the IP10 value each preparation was normalized with that of SMB-PS cells collected 24 h. Y-axis represents the numerical number. Graphical data denote mean + SEM (n=3). **B.** Representative images of transwell chemotaxis tests of SMB-PS and SMB-S15 cells before treatment of resveratrol. BV2 cells on the membrane were stained with crystal violet. **C.** Western blot for PrP^Sc^ in SMB-S15 cells collected at 1, 3 and 7 days after treatment of resveratrol. The cell lysates were digested with 20 mg/ml PK prior to SDS-PAGE. **D.** Western blot for CXCR3 in the lysates of SMB-S15 cells treated with resveratrol for 1, 3 and 7 days. **E.** Measurement of IP10 in the cytolysis and medium of SMB-S15 with ELISA after treatment of resveratrol. Cells were collected 1, 3 and 7 days after exposed to resveratrol or DMSO. DMEM presents the mock cells maintained in culture medium without treatment of resveratrol or DMSO. The IP10 value each preparation was normalized with that of SMB-PS cells treated with resveratrol for 1 day. Y-axis represents the numerical number. Graphical data denote mean + SEM (n=3).

To assess the possible influence of removal of prion propagation on the expression of IP10, SMB-S15 cells were exposed to 10 μM resveratrol for 1, 3, 7 days and the fractions of medium and cells were separately collected. PrP-specific Western blots of the cell lysates revealed clear PK-resistant PrP (PrP^res^) signals in SMB-S15 cells exposed to resveratrol for 1 day, weaker in the preparation of 3 days, and markedly weaker PrP^res^ in the that of 7 days (Fig. 8C). Measurement of IP10 values with ELISA kit in the fractions of medium and cytolysis of SMB-S15 cells identified significantly decreased in cytolysis after treatment of resveratrol for 7 days, and remarkably increased in the medium since exposure to resveratrol for 1 day, in comparison with the cells treated DMSO and mock (Fig. 8D). CXCR3-specific Western blots revealed notably reduction of CXCR3 intensity in SMB-S15 cells treated with resveratrol for 7 days (Fig. 8E). Those data suggest that removal of prion replication in SMB-S15 cells by resveratrol converts the accumulation and secretion of cellular IP10 and recovered the elevation of CXCR3.

## Discussion

Based on our previous study, we have further verified the increase of brain IP10 in several scrapie infected experimental rodent models with different techniques. The increased IP10 colocalizes closely with neurons and microglia but not with astrocytes. Increased IP10 is also identified in a prion infected cell line SMB-S15. More importantly, we have also found an obviously increased CXCR3, the receptor for IP10, both in the brains of scrapie infected animals and in the prion infected cell line.

The increased IP10 and CXCR3 colocalized closely with PrP^Sc^ in the neuronal cells of the brains infected with scrapie agents morphologically, meanwhile, the distributions of those three elements show highly coincidental in the brain regions. Molecular interactions of PrP^Sc^ with IP10 and CXCR3 are verified. Moreover, we have found that more portions of IP10 accumulated insides of the prion infected cell line. Removal of prion propagation in SMB-S15 cells with resveratrol can sufficiently covert IP10 accumulation in cytoplasm and decrease the levels of CXCR3. Those novel data provide a new scenario that chemokine IP10 and its receptor CXCR3 actively participate in prion pathophysiology.

Neuroinflammation is one of the consequences of host innate immunity to various pathological situations in CNS, such as infection, injury, proliferative and degenerative disorders. Meanwhile, neuroinflammation induce subsequently pathophysiological effects, either neuroprotective or neurotoxic. Active neuroinflammation responses have been well documented in prion diseases, represented by activation of microglia, complement system and numerous inflammatory cytokines [26–31]. IP10 is mainly secreted by microglia in CNS after LPS stimulation [32]. Previous studies display the pathological effects of increased IP10 on neuronal cells in different abnormal situations [33–35]. Increased IP10 in brains, CSF and peripheral blood samples of AD cases and AD transgenic mice is repeatedly described [36–38]. In PD patients, the cognitive state and ability are closely related to IP10 levels [39], and increased IP10 promotes the whole inflammatory response and accelerates neuronal cells deaths [40]. The increased brain IP10 during prion infection from current and previous studies display the similar alteration of prion disease as AD and PD.

We have also provided, probably for the first time, along with the increase of brain IP10, its receptor CXCR3 is also upregulated in the brains of scrapie infected mouse models. As the receptor for IP10, monokine induced interferon-gamma (Mig/CXCL9), and interferon-gamma inducible T cell alpha-chemoattractant (I-TAC/CXCL11). CXCR3 functions actively in multiple host physiological and pathological programs, e.g., integrin activation, cytoskeletal changes, suppression of angiogenesis or chemotactic migration, as well as in acute and chronic inflammation, autoimmune diseases, malignant tumors, etc. [41]. CXCR3 critically modulates the activation of glial cells during cuprizone-induced toxic demyelination [42]. The elevated expression of CXCR3 in brain tissues has been documented in many neuroinflammatory and neurodegenerative diseases, such as AD, multiple sclerosis (MS), glioma, chronic pain, human T-lymphotropic virus type 1-associated myelopathy/tropical spastic paraparesis (HAM/TSP) and bipolar disorder [43]. In the postmortem brain tissues of AD cases, CXCR3 is constitutively expressed on a subpopulation of neurons in various regions, including neocortex, hippocampal formation, striatum, cerebellum and spinal cord [44]. The expressions of IP10, Mig and CXCR3 has been observed also in the CNS tissues of MS patients [41]. Obviously, prion disease undergoes similar activation of IP10/CXCR3 signaling in CNS tissues as other neurodegenerative diseases.

Our IHC data here have demonstrated a close morphological association between deposit of PrP^Sc^ and accumulations of IP10 and CXCR3 in brain tissues of prion infected mice. Such histopathological characteristics reflects from not only the colocalizations of those three factors with serial brain sections, but also the distributions in various brain regions with whole brain sections. In AD patients, IP10 positive astrocytes seem to be associated with amyloid deposits [44]. Increased IP10 is intensively colocalized with Aβ-plaques in an AD mouse model [36]. In the demyelinating lesions of postmortem CNS tissue from MS cases, predominantly expressed IP10 and CXCR3 are within the plaques [41]. Areas of plaque formation in MS lesions are infiltrated by CCR5-expressing and CXCR3-expressing cells [45]. The increased IP10 and CXCR3 are thought to be a chronic inflammatory response to the amyloid deposits in AD and MS, and facilize the plaque formation [43]. Increased IP10 and CXCR3 has been also found in the lesions of other diseases with plaque formation, such as atherosclerosis [46, 47] and psoriasis [48]. The close association of increases of IP10 and its receptor CXCR3 with the amyloid deposits of abnormal proteins in various disorders, including prion disease, reflects a similar responsive pattern of the host innate immunity.

Removal of prion propagation in the cell model in this study attenuates the elevated cellular levels of IP10 and CXCR3, providing the evidence that the PrP^Sc^ deposit directly induces the increased IP10 and CXCR3. On the other hand, deprivation of CXCR3 expression may also affect the AD pathology. In an AD transgenic mouse model of amyloid precursor protein (APP)/presenilin 1 (PS1), deletion of CXCR3 significantly reduces the amyloid deposit and Aβ level, facilitates the microglial uptake of Aβ, and improve the spatial memory of the mice [49]. Another study with CXCR3-deficient mice has demonstrated that the CXCR3 deficiency decreases the activation and accumulation of microglial and astrocytes, whilst the mice appear milder clinical course of demyelination during cuprizone feeding and remarkably rapid recovery of body weight after offset of diet [42]. The therapeutic potentials of CXCR3 with different antagonists and monoclonal antibodies for IP10 have been repeatedly evaluated for AD and MS, despite that the mechanism is still not fully clear [43, 50]. Further study of depriving IP10 and/or CXCR3 on prion infection is deserved to explore its therapeutic potential for prion disease.

We have also observed that more portion of the increased IP10 accumulates in the cytoplasm fraction of prion infected cell line. Such pattern can be converted after removal of prion replication. The exact reason is not clear. One speculation is based on active molecular interaction between PrP^Sc^ and IP10. Possibly, more PrP^Sc^ molecules accumulated insides of the prion infected cells will fix more IP10 molecules. After removal of prion agent, the bound-IP10 will be secreted out of the cells. The pathophysiological effects of activation of IP10-CXCR3 signaling in prion infected cells is unknown. An early study has verified that exposures of the human fetal brain cell cultures to simian human immunodeficiency virus (sHIV) sHIV89.6P and viral gp120 cause upregulation of IP10 in neuron, and further induce membrane permeability followed by apoptosis via activation of caspase-3 [33]. Treatment of exogenous IP10 onto cultured neurons results in cell apoptosis via elevating intracellular Ca^2+^ and caspases [51]. Apoptosis and intracellular Ca^2+^ dysfunction are common phenomena in prion pathology, which can be triggered by different pathways [52, 53]. It is reasonable to assume that activation of IP10/CXCR3 signaling in prion infected cells and brains may also involve in such abnormalities.

## Materials and Methods

### Ethical statement

All procedures of animal experiment and housing were performed in accord with the Chinese Regulations for the Administration of Affairs Concerning Experimental Animals. All procedures were approved and supervised by the Ethical Committee of the National Institute for Viral Disease Control and Prevention (IVDC), China CDC.

### Brain samples of scrapie-infected mice

The brain samples of C57BL/6 mice inoculated intracerebrally with mouse-adapted scrapie strains 139A and ME7 were enrolled in this study. The clinical, neuropathological and pathogenic features of these infected mice were described elsewhere [54, 55]. The average incubation times of 139A-, and ME7-infected mice were 183.9±23.1 and 184.2±11.8 days, respectively. Age matched healthy mice were used as control.

### Preparation of brain homogenates

Brain homogenates were prepared based on the protocol described previously [55]. Brain tissues from the infected mice and controls were washed in TBS (10 mM Tris-HCl, 133 mM NaCl, pH 7.4) for three times, and then 10% (w/v) brain homogenates were prepared in cold lysis buffer (100 mM NaCl, 10 mM EDTA, 0.5% Nonidet P-40, 0.5% sodium deoxycholate, 10 mM Tris, pH 7.5) with protease inhibitor Cocktail set III (Merck, 535140, Germany). The tissue debris was removed with low-speed centrifugation at 2000×g for 10 min and the supernatants were collected for further study.

### Cell culture

The cell line SMB-S15 and its normal control cell line SMB-PS were provided by Roslin institute, UK [56]. Cell line SMB-S15 was originally obtained by culture from the brains of mice clinically exposed to scrapie agent Chandler and derived from mesodermal, in which PrP^Sc^ can replicate consistently by cell passage. Cell line SMB-PS was obtained from the SMB-S15 cells treated with pentosane sulfate (PS) without detectable PrP^Sc^ [57]. All cells were cultured in Dulbecco’s modified Eagle’s medium (DMEM) with 10% fetal bovine serum (FBS) at 33 °C humidified atmosphere with 5% CO_2_. The confluent cells were passaged using trypsin/EDTA (0.05 % porcine trypsin and 0.02 % EDTA) every 7 to 10 days, and medium was changed every 3 to 4 days.

### Cell lysates

After washed with PBS for 3 times, the cultured cells were scrapped and harvested by centrifugation at 500×g for 10 min. The pellets were lysed for 1 h using cell protein extraction reagent (CW BioTech, CW0889, China) with protease inhibitor Cocktail set III (1 %, v/v, Merck, 535140, Germany), followed by centrifugation at 500×g for 10 min and the supernatants were collected. Protein concentration was estimated using BCA protein assay kit (Merck, 71285-3, Germany).

### Western blot

Aliquots of protein samples were separated on 12% polyacrylamide gels in Tris-glycine-SDS buffer (SDS-PAGE) and electronically transferred to nitrocellulose membranes (GE, 10600001, USA) using a semi-dry blotting system (Bio-Rad, USA). Membranes were blocked at room temperature (RT) for 1 h with 5 % (w/v) non-fat milk powder in 1×Tris-buffered saline containing 0.1 % Tween 20 (TBST) and then blotted at 4 °C overnight with 1:2000 diluted rabbit anti-CXCR3 (Proteintech, 26756-1-AP, USA) and 1:1000 diluted rabbit anti-IP10 (Abcam, ab9938, UK). After washing with TBST by 3 times, blots were incubated with the corresponding horseradish peroxidase (HRP)-conjugated secondary antibodies at RT for 1 h. The blots were developed using enhanced chemiluminescence system (ECL, PerkinElmer, NEL103E001EA, USA) and visualized on autoradiography films (Carestream, 6535876, China). Images were captured by ChemiDoc™ XRS+ System with Image Lab software (Bio-Rad, USA) and quantified by Image J software.

### Quantitative real-time PCR (qRT-PCR)

Total brain RNAs of healthy and scrapie-infected mice were extracted using the TRIzol (Invitrogen, 15596026, USA) reagent according to the manufacturer’s instruction, followed by degradation of both double- and single-stranded DNA from the samples with RNase-Free DNase (Invitrogen, 2028724, USA). Reverse transcription was performed using SuperScript^TM^ III First-Strand Synthesis System (Invitrogen, 2028724, USA). Briefly, 2 μg of total RNA was mixed with 10 μl RT Reaction Mix (2×), 2 μl RT Enzyme Mix and DEPC-treated water to a volume of 20 μl. The mixtures were maintained at 25°C for 10 min, then incubated at 50°C for 30 min and inactivated by heating at 85 °C for 5 min. To remove RNA from the cDNA, 1μl *E.coli* RNase H was added to the mixture and incubated at 37 °C for 20 min. Aliquots (2 μl) of RT reaction products were amplified by Real-Time PCR. The primers used were designed and synthesized according to the published mRNA sequence (Supplementary Table 1).

Real time-PCR was performed using a supermix containing the fluorescent dye SYBR green (TSINGKE, TSE202, China). Briefly, 1 μl of reverse transcription product, 10 μl of T5 Fast qPCR Mix (2×), 1 μl of each primer (10 μM) and 7 μl of Nuclease-free H_2_O were mixed in a total volume of 20 μl. PCR was performed on a CFX96 Real-Time PCR System (Bio-Rad, USA) under following conditions: after enzyme activation at 40 PCR cycles of 95 °C for 15s, and 60 °C for 60s. The increases in fluorescence were collected and the expressive level of each specific mRNA was determined relative to that of the individual β-actin using the comparative C_t_ method (2^-△△Ct^). All real time-PCR reactions were performed in triplicate.

### Immunofluorescence assay (IFA)

SMB-S15 and -PS cells were washed with PBS and fixed with 4 % paraformaldehyde at RT for 20 min, followed by treatment with 0.4 % Triton X-100 for 5 min and blocked with PBS containing 5% BSA at RT for 1 h. Then, fixed cells were incubated with primary antibodies, including 1:100 diluted pAb anti-IP10, 1:100 diluted pAb anti-CXCR3, 1:100-diluted mAb anti-PrP (Santa, sc-58581, USA) at 4 ℃ overnight. After washing by PBS with 3 times, fixed cells were incubated with 1:200-diluted Alexa Fluor 568-labeled goat-derived anti-rabbit or Alexa Fluor 488-labeled goat-derived anti-mouse antibodies at RT for 1 h. Further, cells were incubated with 1 μg/ml DAPI at RT for 5 min. The images of the targeting proteins were analyzed by a high content screening system (Operetta Enspire, Perkin Elmer, USA). The integral optical density (IOD) values of each field-specific fluorescence staining were collected by software Operetta Enspire. The IOD values of the specific staining were determined relative to that of DAPI-specific staining.

Brain slices were permeabilized with 0.3 % Triton X-100 for 20 min and blocked with BSA for 1 h. After blocked, sections were incubated with 1:100 diluted pAb anti-IP10, 1:100-diluted mAb anti-GFAP (CST, 3670S, USA), 1:100-diluted pAb anti-Iba1 (Abcam, ab5076, UK), 1:100 diluted mAb anti-NeuN (Merck Millipore, MAB377, Germany) or 1:100 diluted pAb anti-CXCR3 at 4 °C overnight. Slices were washed and incubated with 1:200-diluted (v/v) appropriate secondary antibodies at 37 °C for 1 h. After washing, slices were incubated with 1 μg/ml DAPI (Beyotime, China) at RT for 30 min and sealed. The images of the targeting proteins were analyzed by high solution confocal microscopy (LEICA TCS SP8, Germany). Control results using only the 2^nd^ antibodies were shown in supplementary Fig. 1A.

### Immunohistochemical staining (IHC)

Brain tissue was fixed in 10 % buffered formalin solution and paraffin sections (2 μm in thickness) were prepared routinely. After washed with PBS for three times, tissue slices were quenched with 3 % H_2_O_2_ for 10 min and repaired under high temperature with 1 % sodium citrate solution in microwave for 20 min. After blocking in 5% bovine serum albumin (BSA) at RT for 15 min, the sections were incubated with 1:100-diluted pAb anti-IP10, anti-CXCR3, anti-PrP at 4°C overnight. Subsequently, the sections were incubated with 1:250-diluted HRP-conjugated goat-derived anti-rabbit secondary antibody at 37 °C for 1 h, and visualized by incubation with 3, 3’-diaminobenzidine tetrahydrochloride (DAB). The slices were counterstained with hematoxylin, dehydrated and mounted in permount. Control results using only the 2^nd^ antibodies were shown in supplementary Fig. 1B.

### ELISA for IP10

IP10 level of the whole cell lysate and medium were analyzed by a commercial murine IP-10 precoated ELISA kit (NeoBioscience, EMC121.96, China). Briefly, 10 μl of lysate was mixed with dilution buffer supplied by the manufacturer and added to wells of the antibody precoated plate in duplicate. Subsequently, the plate was incubated at 37 °C for 90 min. After washed each well with 300 μl wash buffer for 5 times, 100 μl of biotin-labeled antibodies were added and incubated at 37 °C for 60 min. After washed, the solution containing HRP-conjugated streptavidin were added and incubated at 37 °C for 30 min. The reactions were developed with addition of 100 μl substrate working solution for 15 min in dark and terminated with the stop solution. The OD_450_ values of reaction was measured by a microplate reader (Thermo Scientific, USA). The concentrations of IP10 were calculated by referring to the corresponding standard curve.

### Co-immunoprecipitation (Co-IP)

Cell lysate was aliquoted and used as input controls. A total of 20 mg of cell lysate was incubated at 4 °C overnight with Dynabeads Protein G treated with 4 µg anti-IP10 pAb or 4µg anti-PrP mAb, respectively. Following washed by phosphate-buffered saline (PBS) containing 0.05 % Tween-20 six times, IP products were eluted with elution buffer (50 mM Glycine, pH 2.8) and subjected to PrP specific Western blot together with the corresponding input and isotype IgG.

### Molecular interaction

Interaction between recombinant IP10 and PrP were analyzed using Octet RED96 (ForteBio, USA). The PrP protein was immobilized with the two-fold serially dilution Abs on the 96-well microplate according to manufacturer’s protocol. IP10 protein was injected into each well and IP10-PrP binding responses were measured. The apparent equilibrium dissociation constants (apparent binding affinity, K_D_) for each IP10-PrP interaction were calculated using Octet^®^ RED96 Software [58].

### Transwell chemotaxis test

SMB-PS and SMB-S15 cells were collected after routine trypsinization. 1 ml (approximately 5×10^5^) of each cell line were placed into the lower chamber of Transwell (Corning, 3421). The upper chamber of Transwell moistened with DMEM were carefully placed to the lower chamber. About 1×10^5^ of BV2 cells were placed into the upper chamber. After incubating at 34°C for 24 h, the upper chamber of Transwell were carefully removed. The excessive BV2 cells on the upper chamber membrane were gently wiped out with a moistened cotton swab. After washed with PBS, the membrane was stained with 0.1% crystal violet, and the cells were counted under a light microscope.

### Statistical Analysis

Statistical analysis was conducted using the SPSS 22.0 statistical package. Quantification of IFA were represented as IOD that is the sum of the reaction intensities of all selected objects in the field of view and were performed by Image J software. All experiments in the present study were conducted at least three times with consistent results. All data were presented as the mean+SEM. The P values for differences between two groups were determined by two-tailed Student’s *t* test. The P values were described in figures or expressed as *** (P <0.001), ** (P <0.01), * (P <0.05) and ns (not significant).

## Acknowledgement

This work was supported by SKLID Development Grant (2019SKLID401, 2019SKLID603, 2016SKLID603), National Natural Science Foundation of China (81772197, 81401670, 81630062).

## Author Contributions

Jia Chen contributed to study design, performed experiments and prepared the manuscript. Chao Hu and Wei Yang assisted with the preparations of experimental materials and data analysis. Dong-Dong Chen and Yue-Zhang Wu assisted with the preparations of the experimental animal samples. Lin Wang assisted with statistical analysis. Cao Chen contributed to study design and Xiao-Ping Dong. corresponding authors, contributed to design, study concept, and final manuscript preparation. All authors read and approved the final manuscript.

